# Constitutive Androstane Receptor induces Ribonucleotide Reductase-M2 expression and maintains hepatocyte ploidy in mice

**DOI:** 10.1101/2025.04.29.651109

**Authors:** Anjana Asokakumar, Bhoomika Mathur, Sandip Chorghade, Anthony Chau, Bea Cronologia, Frances Alencastro, Linda Wheeler, Christopher K. Mathews, David D. Moore, Andrew W. Duncan, Sayeepriyadarshini Anakk

**Author notes:** Co-first authors (Anjana Asokakumar, Bhoomika Mathur).

## Abstract

The nuclear receptor Constitutive Androstane Receptor (CAR/NR1i3) is known for regulating various liver functions, including detoxification, nutrient metabolism, and hepatocyte proliferation. While CAR activation has been previously linked to higher ploidy, the underlying mechanisms are not fully known. Here, we uncover a basal role for CAR in maintaining hepatocyte ploidy, such that CAR deletion increases the number of diploid (2c) hepatocytes with a concomitant reduction in tetraploid (4c) hepatocytes. We demonstrate that CAR controls the *de novo* dNTP synthesis by directly transactivating the Ribonucleotide Reductase-M2 (*RRM2*) gene, which encodes the rate-limiting catalytic subunit of the enzyme, ribonucleotide reductase. Further, we find that the ligand-dependent CAR activation is sufficient to induce several genes involved in the *de novo* dNTP synthesis pathways, resulting in higher hepatic dATP and dTTP levels within the liver. Importantly, overexpressing RRM2 levels in CAR knockouts led to increased DNA synthesis and tetraploid (4c) hepatocytes compared to the control mice. Importantly, we demonstrate that CAR-mediated DNA synthesis in liver cells is dependent on catalytically active RRM2 function. Taken together, these findings reveal that the CAR-mediated RRM2 regulation contributes towards DNA synthesis and thereby maintains hepatocyte ploidy.

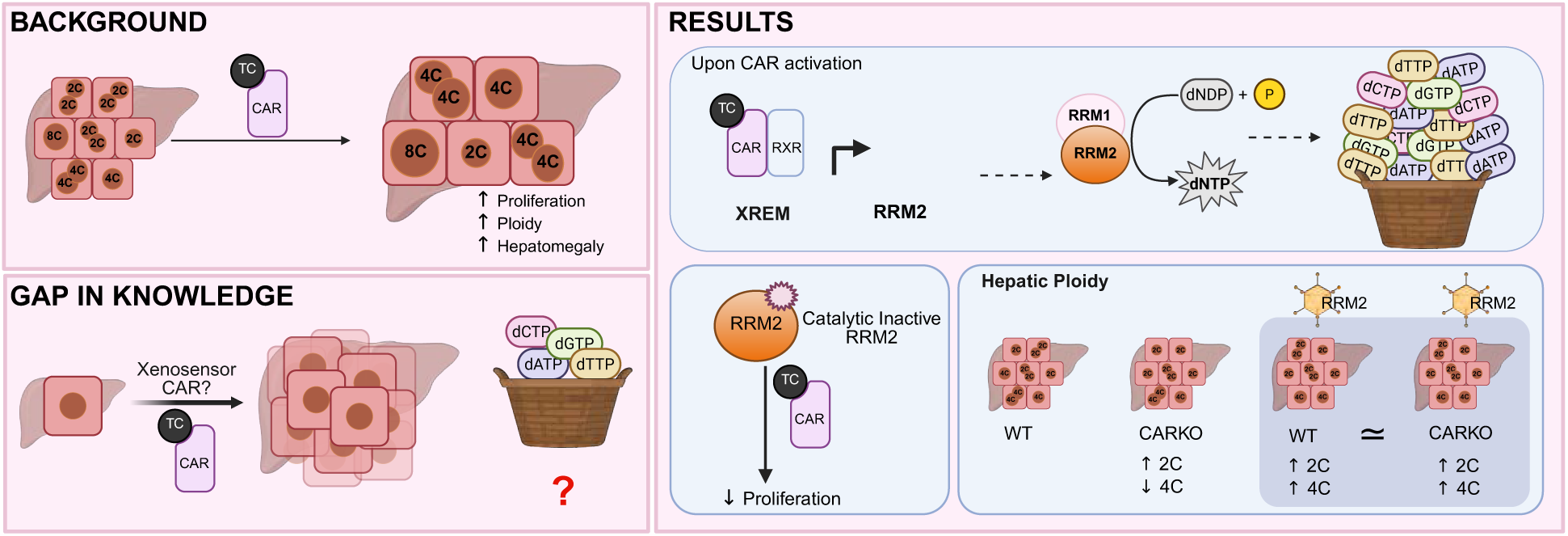

## Introduction

The liver is a metabolic hub and is responsible for many cellular functions(1, 2). Hepatocytes carry out detoxification and intermediary metabolism and comprise 70-80% of the liver tissue(3–5). A salient feature of the hepatocytes is that they exhibit polyploidy(6). Immature developing hepatocytes during the postnatal period (<21 days) are preferentially diploid, while the mature hepatocytes are more polyploid(6, 7). In rodents, polyploidization starts around the time of weaning, resulting in 80-90% of adult hepatocytes being polyploid(6). Some studies have indicated a higher proliferation rate for diploid hepatocytes than polyploid hepatocytes(8, 9). Contrastingly, other studies demonstrate that higher hepatocyte polyploidy supports the adaptations during chronic liver injury or cancer by providing extra gene copies(6–8). Nonetheless, polyploidy is conserved in the liver, but the mechanisms that link some of the metabolic functions, such as detoxification, to polyploidy remain elusive.

A key xenosensor in the liver that transcriptionally regulates detox machinery is the Constitutive Androstane Receptor (CAR or Nr1i3)(10). Upon activation, CAR translocates to the nucleus, heterodimerizes with the Retinoid X receptor (RXR), and binds the xenobiotic response element module (XREM) of the target genes(11–13). Besides detoxification, ligand-mediated CAR activation is typically accompanied by hepatomegaly and increased DNA synthesis as measured by Ki-67(14, 15). Moreover, synthetic CAR activators, TCPOBOP (TC), and phenobarbital have been shown to increase hepatocyte ploidy(14). However, how these cells respond to these CAR-mediated increases in DNA synthesis and increases in dNTP requirements has not been fully examined.

We mined the publicly available transcriptome after CAR activation and found a robust increase in *RRM2* (Ribonucleotide Reductase Regulatory subunit M2) transcripts. This gene encodes the catalytic subunit of the rate-limiting enzyme ribonucleotide reductase, which converts NDPs (Nucleoside diphosphates) to dNDPs (deoxynucleoside diphosphates) in the dNTP synthesis pathway(16–18). Then, we investigated if CAR could bind and activate RRM2 transcription and scanned the 5’ upstream region of the RRM2 gene. We performed luciferase and chromatin immunoprecipitation assays, followed by PCR to validate the functionality of the putative CAR binding site. We measured hepatic dNTP levels to corroborate our findings. We then tested if ectopic expression of RRM2 in CAR knockouts (CARKO) alter proliferation or ploidy status. Finally, we investigated if active RRM2 was necessary for facilitating CAR-mediated DNA synthesis.

## Method

### Mice experiment

C57BL/6J mice were obtained from Jackson Laboratories, and CARKO mice were obtained from Dr Moore’s lab at Baylor College of Medicine. The age and sex of the mice used for the experiments are mentioned in the figure legends. The mice were housed in a standard 12-hr light/dark cycle with chow food and water available ad libitum. All the mice used in the study have complied with NIH guidelines for the care and use of laboratory animals, and experiments were performed as directed by the Institutional Animal Care and Use Committee at the University of Illinois, Urbana-Champaign. All mice were acclimatized to the facility at least 2 weeks prior to the experiments. Mouse CAR agonist, 1,4-bis[2-(3,5-dichloropyridyloxy)] benzene (TCPOBOP, Sigma, #T1443), TC was dissolved in Cornoil and intraperitoneally (IP) injected at 100 mg/Kg once into the mice, euthanized after 3 days of treatment, and collected samples for further studies. Mouse Rrm2 was overexpressed using AAV-DJ8 under the CAG promoter with sfGFP at the C terminus (VB220927-1597gxb and VB220823-1142mex). Viruses were obtained from the vector builder and were IP-injected at a concentration of 10^11 GC. The mice were euthanized after two weeks for serum and liver collection.

### Cell culture

AML12 cells were transfected with 1µg of pCMV-HA–mutRRM2 (mouse; Y176F), pCMV-Myc–WT RRM2 (mouse), and pCMV-Flag–CAR (mouse) using Lipofectamine 3000 for 48 hr. For cotransfection, equal concentrations of individual plasmids were mixed to achieve a total of 1 µg/well. Cells were serum-starved for 16 hr, reseeded at 40% confluence in complete medium containing 10% serum with or without TC (250 nM) dissolved in DMSO, and incubated for 16 hr before analysis. Aliquots of cells were collected at the time of reseeding for immunoblot analysis.

Primary hepatocytes were isolated from WT mice using collagenase perfusion and were treated with hydroxyurea (1mM) or TC - CAR ligand (250nM) or both for 6 hours, and cells were harvested for RNA preparation and qPCR analysis.

### Molecular cloning

Mouse and human CAR open reading frames (ORFs), including wild-type (wtRRM2) and tyrosine-to-phenylalanine substitution mutants (mouse Y176F; human Y175F) (mutRRM2), were synthesized (Twist Bioscience) and cloned into the BamHI–HindIII restriction sites of the pcDNA3 expression vector(pCMV).

### Click-IT Plus Edu Staining

Cell proliferation was assessed by incorporation of 5-ethynyl-2′-deoxyuridine (EdU) during a 2-hr pulse(19). Cells were fixed with 4% paraformaldehyde, and EdU detection was performed using click chemistry according to the manufacturer’s instructions (Thermo Fisher Scientific, # C10640).

### RNA extraction, cDNA synthesis, and qPCR

RNA was isolated from the liver using TRIzol (Invitrogen) according to the manufacturer’s instructions. The quality of the RNA was measured using the Bioanalyzer and 0.2% bleach gel. 3-5 µg of isolated RNA was used to synthesize the cDNA with random primers (NEBiolabs) and the Maxima Reverse transcriptase kit (Thermo Fisher Scientific). 50 ng of cDNA was used for every SYBR green kit (Thermo Fisher Scientific) based qRT-PCR. Relative gene expression was calculated by the ΔCt or ΔΔCt method and normalized to 36b4 or actin, as mentioned in the figure legends. qPCR primer details are listed in the supplementary table.

### Luciferase assay

HepG2 cells in a 6-well plate were transfected using calcium phosphate precipitation as described in previous publications(20, 21). 0.1µg/well of luciferase reporter (mRRM2-pTKLUC or mRRM2mutant-pTKLUC), 0.05µg/well of pCMX-CAR, 5ng/well of RXR, and 0.1µg/well of β-gal were used for the luciferase assay. PGEM4 was used to scale the total plasmid concentration to 5µg/well. 35 bp of the CAR response element identified in the RRM2 promoter region was sequenced and cloned into TK-LUC for analysis. The mutant RRM2 CAR response element was generated by mutating the key GG nucleotides to AA. After transfection, the cells were treated with TC: 250 nm overnight. Luciferase expression was assayed and normalized using β-gal expression. All luciferase values were averaged from triplicate samples. All transfection data were repeated at least three times.

### Western blot assay

Protein lysates were made with RIPA buffer (25 mM Tris-HCl, 150 mM NaCl, 0.5% Sodium deoxycholate, 1% Triton X-100, 0.1% SDS, 0.1 mL Phosphatase inhibitor, and 1x protein inhibitor). A total of 25µg of liver tissue lysate or 30µg of cell lysate was loaded to each lane and was separated with 10% SDS-PAGE. The proteins were then transferred to a nitrocellulose membrane. After blocking with 5% non-fat milk in TBS, the membrane was probed with a primary antibody overnight at 4 ^0^C (Gapdh (# 2118, Cell Signaling, 1:5000 dilution), RRM1 (# sc-11733, Santa Cruz, 1:100 dilution), RRM2 (# sc-10848, Santa Cruz, 1:100 dilution), RRM2 (# sc-398294, Santa Cruz, 1:100 dilution), Flag (# F1804, Sigma, 1:1000 dilution), HA (# 51064-2-AP, Proteintech, 1:5000 dilution), Myc (# sc-40, Santa Cruz, 1:1000 dilution), His (# ab18184, Abcam, 1:1000 dilution) followed by secondary antibody 1 hr incubation at RT (anti-mouse # 115-005-174, anti-rabbit # 111-005-045, anti-goat # 805-005-180 from Jackson Immunoresearch at 1:5000 dilution). The membrane was developed with an enhanced chemiluminescent detection reagent and imaged with X-ray film or a Bio-Rad ChemiDoc.

### Liver histology and staining

Mouse liver tissues were fixed in 10% formalin at 4 ^0^C for 12-24 hours, processed, and embedded in paraffin. 5 µm liver sections were made and deparaffinized in xylene and hydrated in a series of graded alcohol and water. Standard hematoxylin and eosin (H&E) staining was used to examine morphology. For immunohistochemistry (IHC), sections were incubated with 3% hydrogen peroxide for 15 minutes to block endogenous peroxidase. In both IHC and immunofluorescence (IF), the sections are heated in Tris-EDTA (pH 9.0) for 30 minutes for antigen retrieval, followed by 1 hour of blocking in 5% BSA and 2% NGS at 4 ^0^C. The sections were incubated with the primary antibody Ki-67 (# 550609, BD Bioscience, 1:100 dilution), PCNA (# sc-25280, Santa Cruz, 1:750 dilution), pHH3 # PA5-17869, Invitrogen, 1:750 dilution), β-catenin (BDB610154, BD Bioscience, 1:1000 dilution), or H2A.X (ab11175, abcam, 1:500) at 4 ^0^C overnight. After the TBS-0.025% Triton X-100 wash, samples were incubated with horseradish peroxidase (HRP) anti-mouse (# 115-005-003, Jackson ImmunoResearch, 1:750 dilution) or anti-rabbit # 111-005-045, Jackson ImmunoResearch, 1:750 dilution) or anti-mouse 555 (# A-21424, Invitrogen, 1:1000 dilution) or anti-rabbit 555 (# A-21428, Invitrogen, 1:1000 dilution) fluorophore for 1 hour at RT. For IHC, the DAB staining kit (Vector Laboratories, #SK-4100) was used according to the manufacturer’s protocol. For IF, the sections were incubated with NuBlue (Thermo Scientific, #R37605) for 30 minutes at room temperature, and the background signals were quenched using Vector TrueVIEW Autofluorescence Quenching (#SP-8400-15) from Vector Laboratories as per the instructions. The slides were viewed with Nanozoomer or the LSM 900 confocal microscope.

### Tunnel staining

For checking the DNA damage, tunnel staining was performed with Click-iT™ Plus TUNEL Assay Kits (ThermoFisher Scientific, # C10617) as per the instructor’s manual and viewed under the LSM 900 microscope.

### Chromatin Immuno-Precipitation (ChIP)

Briefly, the FLAG-CAR construct was transiently transfected into HepG2 cells. After 24 hours of TC treatment, the FLAG-tagged CAR was pulled down using an anti-FLAG antibody (Sigma, # F1840), followed by sonication and PCR amplification of the RRM2 promoter region. 5’CTCCCAAAGTGCTGGGATTA3’ and 5’CAAGCGACCAGGCTTCTTAC3’ primers were used to for human RRM2 promoter DNA fragments. All transfection and pull-down experiments were performed in three separate wells, and IgG was used as a control.

### dNTP Pool analysis

∼1g of liver tissue was washed with ice-cold 10mM Tris buffer, pH 7.3. 5x volume of ice-cold Tris buffer relative to the weight of the tissue was added to the minced tissue, and the mixture was homogenized completely. Cell debris was removed by centrifugation at 100g for 10 minutes. Supernatant was transferred for further analysis and protein measurement. 100% ice-cold methanol was added to the supernatant to obtain a final concentration of 60% methanol (v/v), followed by incubation at 20 ^0^C for 1 hr with intermediate vortexing. The samples were then heated in boiling water for 3 minutes and centrifuged at 17,000 g for 15 minutes at 4 ^0^C. The resulting supernatant was transferred to a fresh tube and completely dried in a speed vacuum or lyophilizer. The dried extracts were resuspended in water, and dNTP concentrations were determined as previously described(22, 23).

### ChIP-seq data analysis

We analyzed the BigWig and BED files (GSE112199) for YFP-tagged mouse and human CAR ChIP-seq from Niu et al(24). Tracks were viewed on IGV aligned to mm10 assembly.

### RNA-seq data analysis

We analyzed the processed RNA sequencing file from WT mouse liver treated with Cornoil or TC (GSE98666) by Cheng et al(25).

### Hepatocyte isolation

Hepatocytes were isolated as described in Wilkinson et al(8).

### Ploidy analysis

For detection of cellular ploidy, freshly isolated primary hepatocytes were stained with 2 μl/ml Fixable Viability Dye (FVD) eFluor 780 (eBioscience, San Diego, CA) and 15 μg/ml Hoechst 33342 (Life Technologies, Carlsbad, CA), as described(8). For nuclear ploidy, nuclei from the frozen liver tissue were isolated as previously described(9, 26). - A piece of frozen liver tissue, approximately 5 x 5 x 3 mm, was thawed at room temperature in 1 mL of PBS and chopped into tiny pieces with a scalpel blade. The liver suspension was passed through a 20-gauge needle (10 times) and a 23-gauge needle (10 times) to break open cells. Suspensions were fixed overnight in 70% ethanol at 4°C, washed with PBS, centrifuged at 500g for 5 minutes at 4°C, and digested in 1.5mL pepsin (0.5mg/mL in 0.1N HCl, Millipore Sigma Life Science) for 20 minutes at RT to generate nuclear suspensions. The nuclear suspensions were washed with 3 mL nuclear wash buffer (PBS, 0.1% BSA, 0.5% Tween-20), incubated with 1.5 mL 2N HCl for 12 minutes at 37°C, washed with 3mL cold borate buffer (6.25g H3BO3/L type I water, pH 8.5), and centrifuged at 500 g for 5 minutes at 4°C. The isolated nuclei were then stained with 5 μg/mL propidium iodide (Invitrogen, Carlsbad, CA) in PBS with 250 μg/mL RNase A (Invitrogen) and analyzed with an LSR II flow cytometer (BD Biosciences, Franklin Lakes, NJ) running BD FACSDiva™ Software pH v9.0. FACS plots were generated using FlowJo 10.8.2 (FlowJo LLC, Ashland, OR)

### Image analysis software

ImageJ was used to measure cellular area and quantify Ki-67, PCNA, and pHH3 staining. QuPath is used to count the total cells.

### Statistical analyses

All statistical analyses were performed using GraphPad PRISM (GraphPad Software, Inc., La Jolla, CA). Data are presented as mean ± standard deviation (SD) or mean ± standard error of the mean (SEM). Two-way ANOVA was performed unless mentioned in the figure legends. Level of significance was \**P* < 0.05, \*\**P* < 0.01, \*\*\**P* < 0.001.

## Results and Discussion

### Loss of CAR alters 2c and 4c ploidy status in the hepatocytes

As expected, CAR activation by TC results in an increased Ki-67 staining, hepatomegaly, and robust induction of its downstream target genes, including *Cyp2b10,* Cyclin A (*CcnA*), and Cyclin B (*CcnB*) in the liver(27, 28), and these responses are absent in the global CAR knockout mice (CARKO) (Fig 1A, Supplementary Fig 1A-G). Besides these known targets, we found that the E2F family of transcription factors (*E2f1* and *E2f8*), responsible for controlling liver polyploidization, are upregulated in a CAR-dependent manner (Fig 1 B-C). Similarly, the transcript levels of Aurora Kinase B (*AurkB*) and Polo-like kinase1 (*Plk1*), genes responsible for the mitotic entry, are increased upon CAR activation(29) (Supplementary Fig 1 H-I). These findings highlight a key role for CAR in regulating hepatic polyploidy. Next, we examined the cellular ploidy of 5-week-old male WT and CARKO hepatocytes and found that deletion of CAR resulted in higher 2c content that correlated with the reduction in 4c content. However, 8c and 16c proportions, which account for ∼20-30% of the polyploid state, were not altered (Fig 1D-F). Taken together, these findings suggest that CAR regulates transcriptional regulators of ploidy and is necessary to maintain 2c and 4c-hepatocyte ratios (∼70-80% of ploidy states) and their distribution.

**Figure 1.**
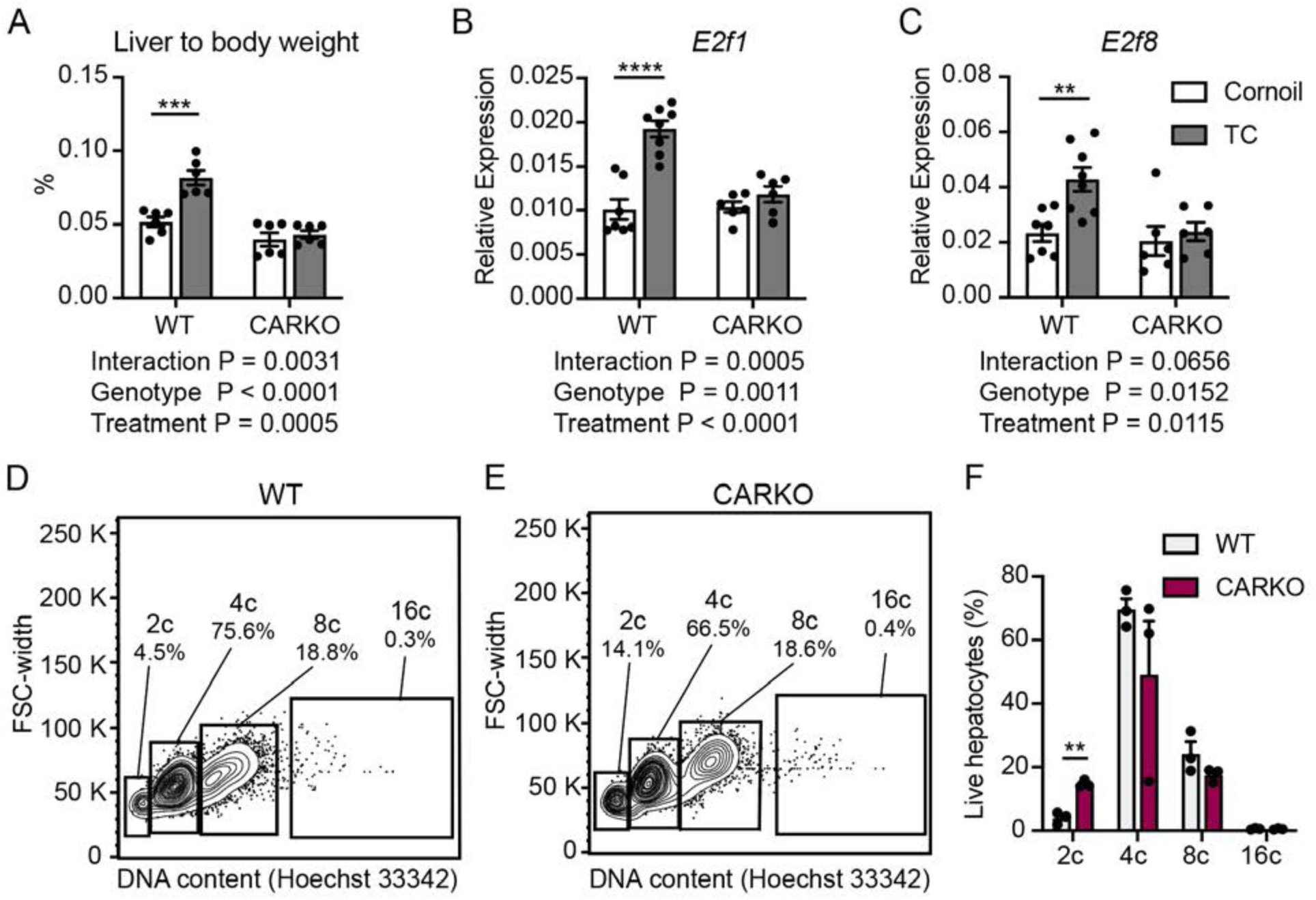
Loss of CAR disrupts the mouse hepatocyte ploidy. (A) The liver-to-body weight ratio of WT and CARKO mice on CAR activation with TC or with control (Cornoil). (B-C) The transcript levels of ploidy-regulating genes *E12f1* and *E2f8* on CAR activation were analyzed using qPCR (n = 6-8 mice/group, adult male mice, 3-month-old; 36b4 was used as the internal control). Two-way ANOVA was performed for the statistical significance. (D and E) Isolated hepatocytes were isolated from 5-week-old male mice, FACS sorted, and used for cellular ploidy analysis. Representative images of gating channels for identifying the DNA content of WT (D) and CARKO (E) hepatocytes. (F) Quantification of live hepatocytes obtained from cellular ploidy analysis and unpaired two-tailed T-tests was used to perform the analysis, **p<0.005, ***p<0.0005.

### CAR regulates the transcription of the *Rrm2* gene

To investigate the mechanisms by which CAR activation promotes DNA synthesis and subsequently ploidy, we mined publicly available CAR transcriptome accessible through the Nuclear Receptor Signaling Website Atlas (NURSA) (http://signalingpathways.org/datasets/dataset.jsf?doi=10.1621/datasets.01003), which looked at post-3 days of TC treatment. Apart from the well-known increase in cyclin gene expression, we noted an unexpected induction of ribonucleotide reductase subunits *Rrm1* and *Rrm2* in response to TC treatment. The *Rrm1* gene encodes the substrate-binding domain, while the *Rrm2* gene encodes the catalytic subunit of the rate-limiting enzyme in the *de novo* dNTP synthesis pathway - Ribonucleotide reductase(17). We validated CAR-mediated induction of *Rrm1* and *Rrm2* transcripts (Fig 2 A-B). Whereas the alternative subunit, *Rrm2b,* induced in response to DNA damage(17, 30) was downregulated (Fig 2C). We also found that only the RRM2 protein was increased upon CAR activation, not RRM1 (Fig 2D-F). These upregulations were absent in the TC-treated CARKO mice, suggesting CAR was necessary for these transcript and protein changes.

**Figure 2.**
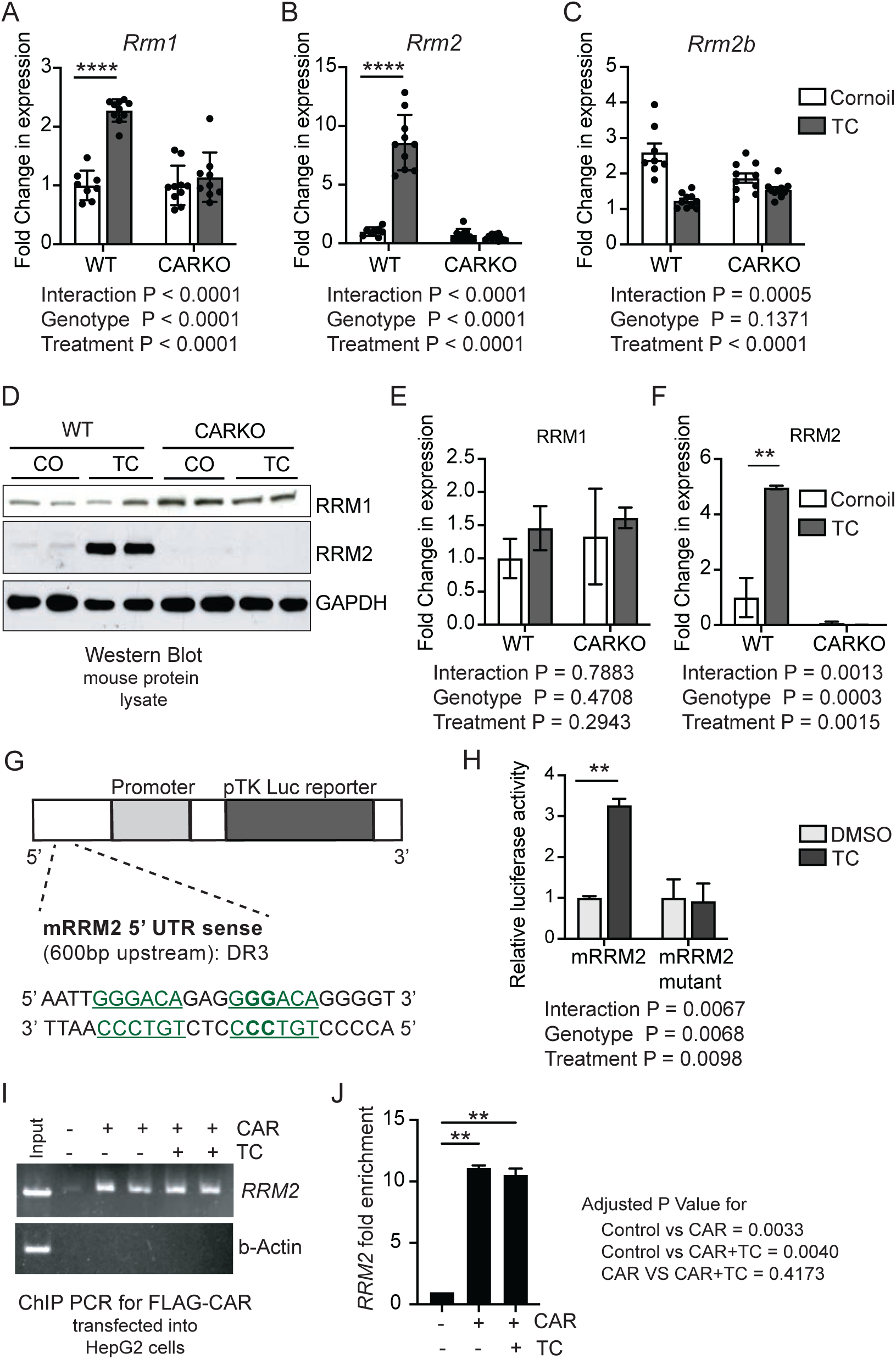
CAR binds and transactivates *RRM2* expression. Transcript expression levels of the ribonucleotide reductase subunit, *Rrm1* (A), and *Rrm2* (B) were analyzed using qPCR. (C) *Rrm2b*, an alternative subunit of ribonucleotide reductase, was also measured by qPCR. (n = 8-10 mice/group, 3-month-old male mice; β-actin was used as the internal control for qPCR analysis) Two-way ANOVA was performed, and graphs are represented as mean ± SEM. (D) Immunoblot analysis of RRM1 and RRM2 in the mouse liver lysates with and without TC treatment and (E-F) normalized quantification are represented as mean ± SD. (G) Schematic representation of the luciferase assay and the target RRM2 site sequence used for the assay. The mutated sites for mRRM2 mutants are marked in **bold** and (H) the quantification represented as mean ± SEM. (I-J) FLAG-tagged CAR construct was transfected into HepG2 cells, and the expressed FLAG-CAR proteins were pulled down to perform ChIP. ChIP-PCR was performed for the analysis (I) and quantified (J), and the graph is represented as mean ± SD (n = 3 technical replicates). Two-way ANOVA was performed on A-C, E-F, and H, while One-way ANOVA was performed on J.

Based on these findings, we examined whether CAR directly regulates *Rrm2* transcription. We screened the *Rrm2* promoter regions and identified putative CAR binding sites. The putative CAR-binding DR3 motif (AGGT/GCA)(28) that was present 600 bp upstream in the *Rrm2* promoter region was then cloned into the pTK-Luc vector (Fig 2G) to test its functionality using a luciferase assay. CAR activation with TC led to a significant increase in luciferase activity, which was lost when this DR3 site on *Rrm2* was mutated (Fig 2H), confirming the transcriptional activation. We then tested if CAR directly binds to the *Rrm2* promoter region using a chromatin immunoprecipitation PCR (ChIP-PCR). FLAG-tagged-CAR-transfected HepG2 cells were treated with TC, and CAR was pulled down using the FLAG antibody. ChIP-PCR analysis showed that ectopic expression of CAR was sufficient to allow its binding to the *Rrm2* gene (Fig 2 I-J), irrespective of the presence or absence of TC. To further validate these findings *in vivo*, we mined the previously published hepatic CAR ChIP-seq data (Supplementary Fig 2), which was performed in CARKO supplemented with either mouse or human CAR, followed by activation with their respective ligands(24). *Cyp2b10* was used as a gold standard downstream CAR target, and as expected, it showed robust CAR binding peaks upon activation with phenobarbital (PB), TC in mouse CAR, and CITCO in the humanized CAR mice (Supplementary Fig 2A). Importantly, we found ligand-induced CAR binding peaks in *Rrm2* were conserved in mice and human (Supplementary Fig 2C). These results demonstrate that the CAR can bind and promote the transcription of the *Rrm2* gene.

### Activation of CAR upregulates the *de novo* dNTP synthesis pathway

We next tested if CAR activation regulated other genes encoding enzymes in the dNTP metabolic pathway besides the rate-limiting catalytic subunit of ribonucleotide reductase, *Rrm2*. To test this, we mined previously published bulk RNA sequencing data after TC treatment(25). Our analysis revealed that apart from the expected increases in cyclins, CAR regulated the entire dNTP pathway from synthesis to salvage, as well as genes involved in ploidy and cytokinesis (Supplementary Fig 3). We first focused on dNTP synthesis and validated many of these changes with qPCR.

dNTP synthesis pathway includes several steps, with ATIC (5-aminoimidazole-4-carboxamide ribonucleotide formyltransferase/IMP cyclohydrolase) and UMP synthase (Uridine monophosphate) being responsible for *de novo* purine and pyrimidine biosynthesis pathways, respectively(31, 32). Briefly, IMPs are converted to ADPs (Adenose diphosphate) or GDPs (Guanosine diphosphate), while UMPs are converted to UDP (Uridine diphosphate) and to UTP (Uridine triphosphate), which is then acted upon by CTP synthetase to generate CTP (Cytidine triphosphate). Ribonucleotide reductase subsequently reduces these substrates to their respective dNDPs (deoxynucleoside diphosphate)(33). dCMP deaminase enzyme keeps the pyrimidine dNTP pool by converting dCMP to dUMP, which in turn can be further modified to dTDP by deoxythymidylate (dTMP) kinase and finally to dNTPs via NDP (nucleoside diphosphate) kinase (Figure 3 A).

**Figure 3.**
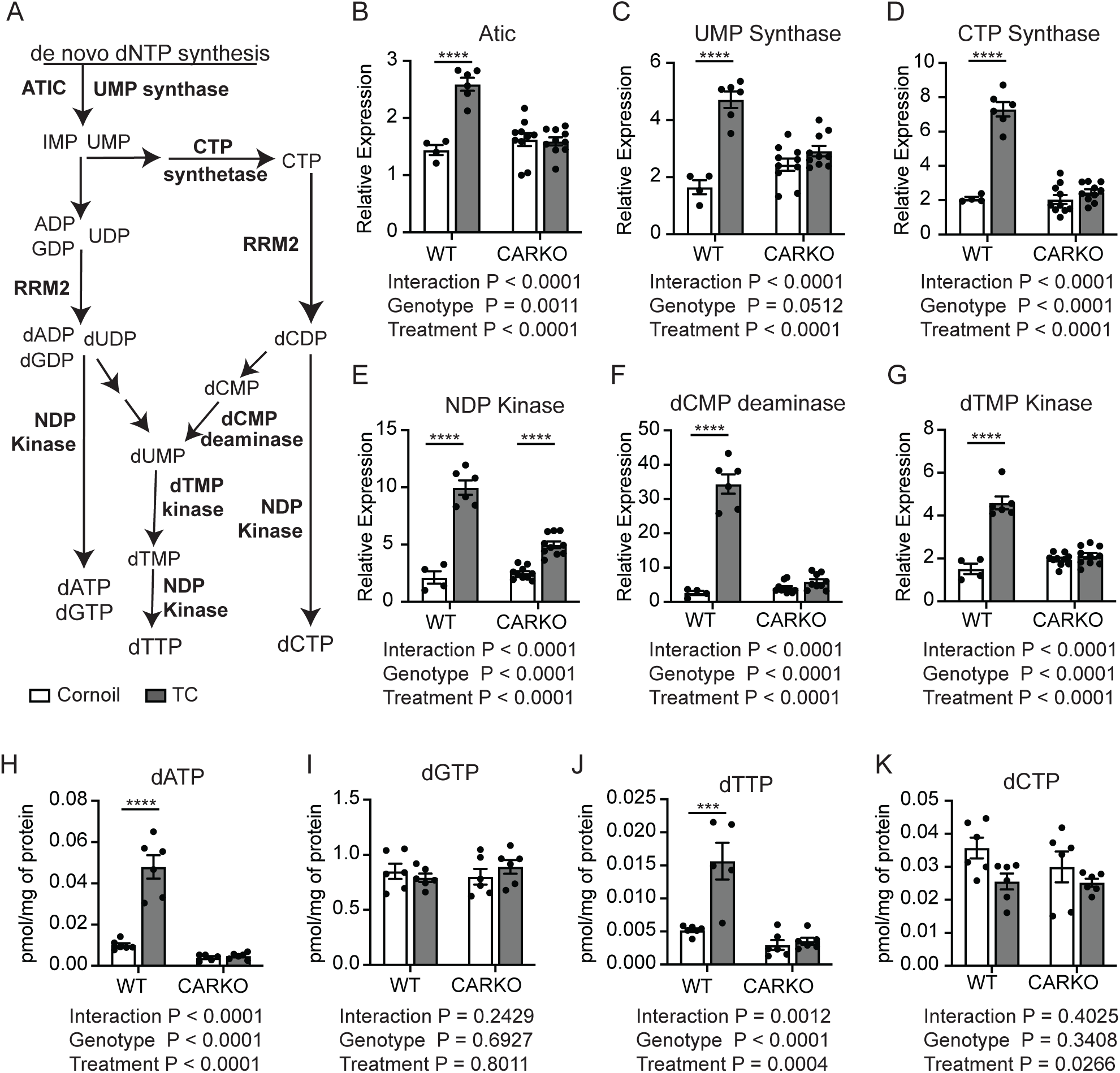
CAR activation increases key enzyme transcript levels in the *de novo* dNTP synthesis pathways. (A) Schematic representation of the crucial enzymes involved in the de novo dNTP synthesis pathway. (B-G) Transcript levels of these enzymes – ATIC (B), UMP Synthase (C), CTP Synthase (D), NDP Kinase (E), dCMP deaminase (F), dTMP Kinase (G) were measured using qPCR post-3 days of TC treatment. (n = 4-6, 3-month-old male mice, b-actin was used as the internal control for qPCR analysis). (H-K) Nucleotide (dATP (H), dGTP (I), dTTP (J), dCTP (K)) levels were analyzed from the liver tissue on CAR activation (n = 4-6 mice/group, 3-month male mice, normalized to the total protein of the liver sample). Two-ANOVA was performed and all the graphs are represented as mean ± SEM.

TC treatment in WT mice resulted in the upregulation of transcripts encoding for these key enzymes – ATIC, UMP synthase, CTP synthetase, NDP kinase, dCMP deaminase, and dTMP kinase in the liver (Figure 3 B-G). Despite the CAR-independent increase in NDP kinase, induction of other enzymes was lost in the CARKO, indicating CAR dependency and highlighting that CAR may regulate the expression of genes in the entire dNTP synthesis pathway. Moreover, ChIP-seq data revealed that several genes (*Rrm2, Atic, NDP kinase encoded by Nme1*) involved in dNTP synthesis exhibited CAR binding peaks (Supplementary Fig 2 C-E), validating our data(24).

Based on these results, we examined the dNTP pools in the liver tissue with and without CAR activation. Consistent with the transcriptional data, dATP and dTTP levels were robustly induced in a CAR-dependent manner (Figure 3 H and J). Intriguingly, the dCTP and dGTP levels were unchanged. One possibility is that the dGTP are present in high concentrations (∼1 pg/mg of protein) in the liver subsequent to enriched mitochondria pool(34) (Figure 3I). Whereas dCTP levels may be converted to dTTP due to increase dCMP deaminase transcript (Figure 3 K).

Another possibility is the dATP is a known allosteric inhibitor of RRM2 activity(35, 36), thus an increase in dATP will in turn inhibit RRM2 levels. Nonetheless, these results uncover that CAR activation upregulates dNTP synthesis genes and certain dNTP pools.

### Ectopic RRM2 expression induces DNA synthesis and alters ploidy in CARKO hepatocytes

Since RRM2 mediates the rate-limiting step for *de novo* dNTP synthesis(37), we examined whether overexpressing (OE) RRM2 in the CARKO mice can compensate for ploidy loss and promote DNA synthesis. Using AAV machinery, we introduced a 10^11 GC virus carrying RRM2-GFP or Control-GFP construct into the mice and ensured that this did not cause overt inflammation (Supplementary Fig 4 A-D). As expected, compared to the control-GFP virus, RRM2-AAV led to a robust increase in *Rrm2* mRNA in WT and CARKO mice, ∼10 fold and 6-fold respectively (Fig 4 A). Importantly, RRM2 overexpression did not change the *Rrm2b* or *Rrm1* transcript levels in both WT and CARKO mice, indicative of specificity of this overexpression with no compensatory impact on other subunits (Fig 4 B, Supplementary Fig 4 E). We validated the RRM2-GFP expression in the livers of WT and CARKO mice by imaging isolated primary hepatocytes. Despite a lower fold increase in *Rrm2* transcript levels in CARKO, we observed no significant change in the RRM2-GFP expression compared to WT. (Fig 4 D)

**Fig 4.**
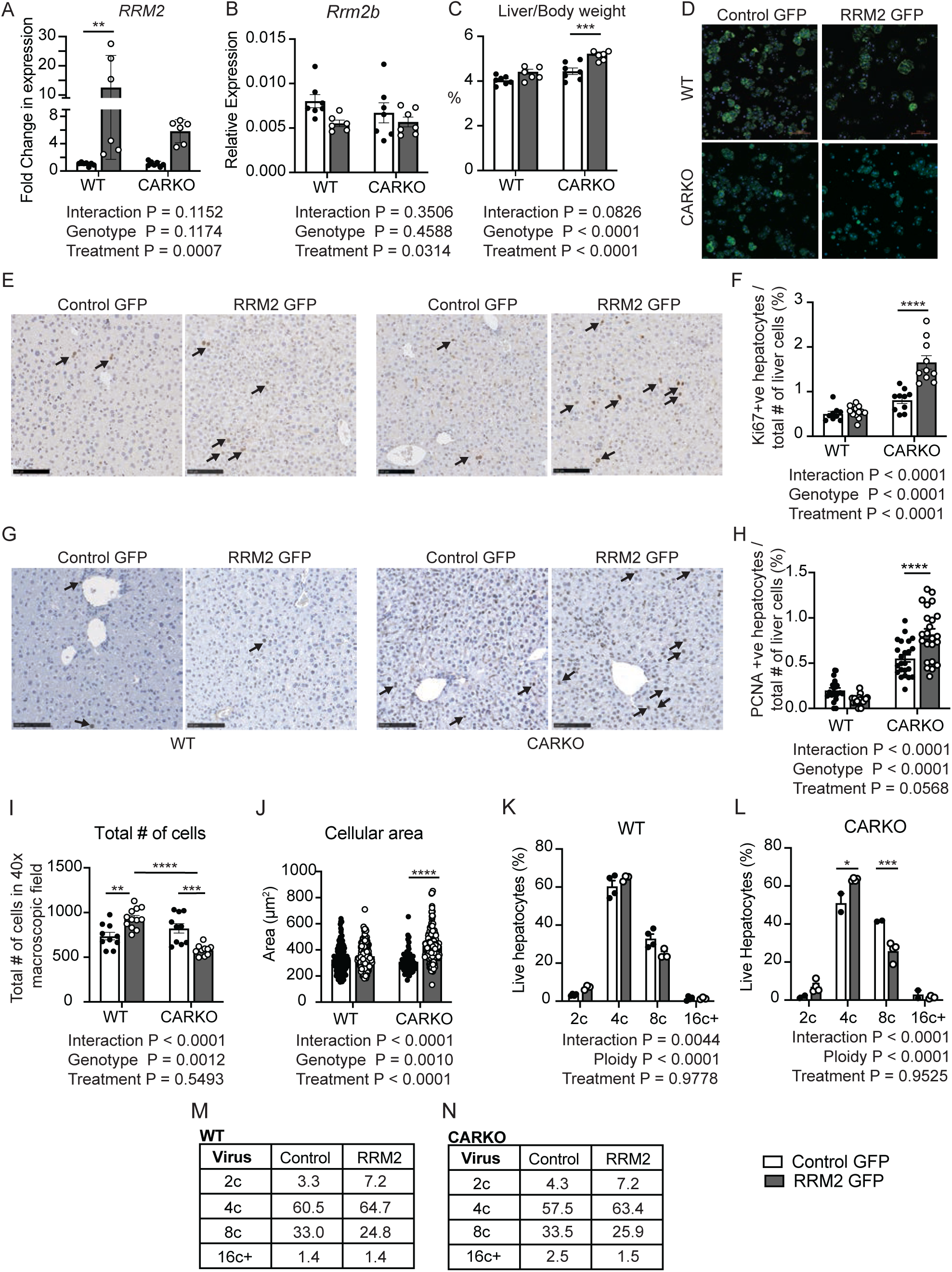
O**v**erexpression **of RRM2 in CARKO increases hepatocyte proliferation and alters cellular ploidy.** Transcript levels of *Rrm2* (A) and *Rrm2b* (B) from WT and CARKO mice livers treated with Control GFP or RRM2-GFP. (n = 6-8 mice/group, 4-month-old female mice, 36b4 was used as the internal control for qPCR analysis). (C) The liver-to-body weight ratio for the virus-treated mice. (D) Hepatocytes from mice treated with control-GFP and RRM2-GFP virus were isolated with a 2-step collagenase perfusion. These cells were incubated with DAPI (blue) and viewed under a microscope for GFP or RRM2-GFP (green) expression. Representative images of (E) Ki-67 and (G) PCNA from WT and CARKO liver treated with control GFP or RRM2-GFP. Black arrows point to the Ki-67 or PCNA staining. Quantification of (F) Ki-67 and (H) PCNA -positive hepatocytes in the liver section was normalized to the total number of cells under the focus region. (I) Total number of cells under the 40x Microscopic field in the liver sections. (J) Liver sections were stained with β-catenin and DAPI, and each cell’s cellular area was measured with ImageJ software. (n = 6-8 mice/group, technical replicate = 3 per group). (K-L) Cellular ploidy quantification for WT and CARKO with control GFP or RRM2-GFP. Hepatocytes were isolated by 2-step collagenase perfusion and proceeded for FACS sorting and ploidy analysis. (M-N) Average of cellular ploidy of WT (M) and CARKO (N) across 2-3 samples per group. Two-way ANOVA was performed for all the analyses, and the graphs are expressed as mean±SEM.

Nonetheless, we were surprised to notice an 18% increase in the liver-to-body weight ratio in CARKOs but not in the WT mice (Fig 4 C, Supplementary Fig 4 F-G). To examine if there was excessive DNA synthesis upon RRM2 OE in CARKO, we used the DNA synthesis marker, Ki-67 and PCNA which stains the entire cell cycle but with high intensity in the S phase(38–40). RRM2 OE dramatically increased the percentage of Ki-67 and PCNA positive nuclei, specifically in CARKO (∼2 fold) compared to WT mice (Fig 4 E-H, Supplementary Fig 5 A-D, Supplementary Fig 6). Intriguingly, despite higher staining of these markers, the total number of cells in the given microscopic field in CARKO was lower (Fig 4 I). Furthermore, when we measured the hepatocyte cellular area on RRM2 OE using β-catenin staining, CARKO exhibited larger hepatocytes compared to WT (Fig 4 J, Supplementary Fig 5 E-H). Overall, these experiments show that introducing RRM2 in CARKO is sufficient to increase DNA synthesis, cellular area, and, subsequently, the hepatosomatic index (liver-to-body weight ratio).

In WT adult mouse livers, approximately 80-90% of hepatocytes are polyploid. Further, hepatocytes can be mono- or binucleate, resulting in varying cellular and nuclear ploidy(7, 41). For instance, a tetraploid can be 4c in a mononucleate or 2c in a binucleate cell; an octoploid can be 8c in a mononucleate or 4c in a binucleate cell, and so on. Hence, the percentage of 2c in nuclear ploidy analysis will be higher than the cellular ploidy. As expected, nuclear ploidy distribution differs from the cellular ploidy shown earlier. In nuclear ploidy, 2c and 4c nuclear content comprises ≥90 percent of all the nuclei. Nuclear ploidy was assessed in frozen liver samples that were also used for transcriptional analysis elsewhere. RRM2 OE in WT mice did not alter nuclear ploidy status in the hepatocytes (Supplementary Fig 7 A, C). However, in the CARKO hepatocytes, RRM2 OE restored the nuclear 2c and 4c to WT levels (Supplementary Fig 7 B-C). This data is consistent, as most tetraploid hepatocytes (70-80%) are binucleate with diploid nuclei, which explains the increase in 2c nuclei in the CARKO RRM2 OE group. Likewise, as most octoploid hepatocytes are binucleate with tetraploid nuclei, this may lead to a decrease in 4c nuclei.

Additionally, in a separate cohort of mice, we also determined cellular ploidy after RRM2 OE. Although RRM2 OE increased the number of 2c hepatocytes in both genotypes, this increase was insignificant (Fig 4 K-N). However, ectopic expression of RRM2 in CARKO increased the percentage of 4c and 8c hepatocytes. Both these changes in the CARKO were comparable to those of WT mice treated with RRM2. Nonetheless, RRM2 OE in CARKO results in higher Ki-67 staining, increased 2c nuclei, and increased 2c hepatocytes.

Unlike 5-week-old mice, whose livers are still maturing, 16-week-old adult livers are in a quiescent state and exhibit a different ploidy distribution. We found that the older adult mice displayed more 8c hepatocytes, which is consistent with a concomitant decrease in 4c hepatocytes (Supplementary Fig 6 D). In the CARKOs, we noticed that 2c hepatocytes significantly dropped with aging from 14.1% to 4.3% (Fig 1 E-F and Fig 4 L, N), indicating that ploidy “catches up” over time.

Since ploidy can be altered with pathological conditions(7, 41), we examined whether this change in the ploidy seen in WT and CARKO post-RRM2 OE was pathological by tunnel and H2Ax staining for DNA damage. We found no immunostaining differences between the control and RRM2 OE, negating the role of pathological ploidy changes; rather, they are driven by RRM2 OE (Supplementary Fig 7 E-F). Collectively, these data suggest that the CAR contributes towards dNTP synthesis and hepatocyte ploidy via RRM2.

### Active RRM2 is necessary for CAR-mediated DNA synthesis

Finally, we tested the dependence of CAR on RRM2 function in orchestrating cellular proliferation. We transiently inhibited RRM2 using hydroxyurea (HU) and tested for TC-mediated CAR activation in primary mouse hepatocytes. As expected, TC induced *Rrm2, Ccnb1,* and dNTP synthesis gene transcripts in cultured WT hepatocytes. Importantly, these upregulations were lost upon co-treatment with HU (Figure 5A), underscoring the relevance of RRM2 in facilitating CAR responses. But this still did not directly assess for DNA synthesis upon CAR activation. Therefore, we performed an EdU incorporation experiment with and without functional RRM2 by generating wtRRM2 and catalytically inactive RRM2(42, 43) (mouse Y176F) constructs (Fig 5 B-D, Supplementary Fig 9). Since the transfection efficiency of multiple plasmids are technically challenging in primary hepatocytes, we utilized AML12, an immortalized non-tumorigenic mouse hepatocyte cell line, for our studies. We also took advantage of the fact that these cells had poor expression of RRM2, consistent with the quiescent hepatocytes under normal physiology. Transfecting these cells with CAR did not alter EdU incorporation, whereas introducing wtRRM2 increased it, which was further amplified by co-transfecting CAR and wtRRM2 (Fig 5 B-C, Supplementary Fig 9 A-B). This increase was completely lost when mutRRM2 (Y176F) was co-transfected with CAR, but the reintroduction of wtRRM2 restored it (Fig 5 B-C, Supplementary Fig 9 A-B). Next, we examined DNA synthesis upon CAR activation under this same experimental paradigm. Activating CAR increased EdU incorporation as expected, and it was dampened in the presence of mutRRM2.

**Fig 5.**
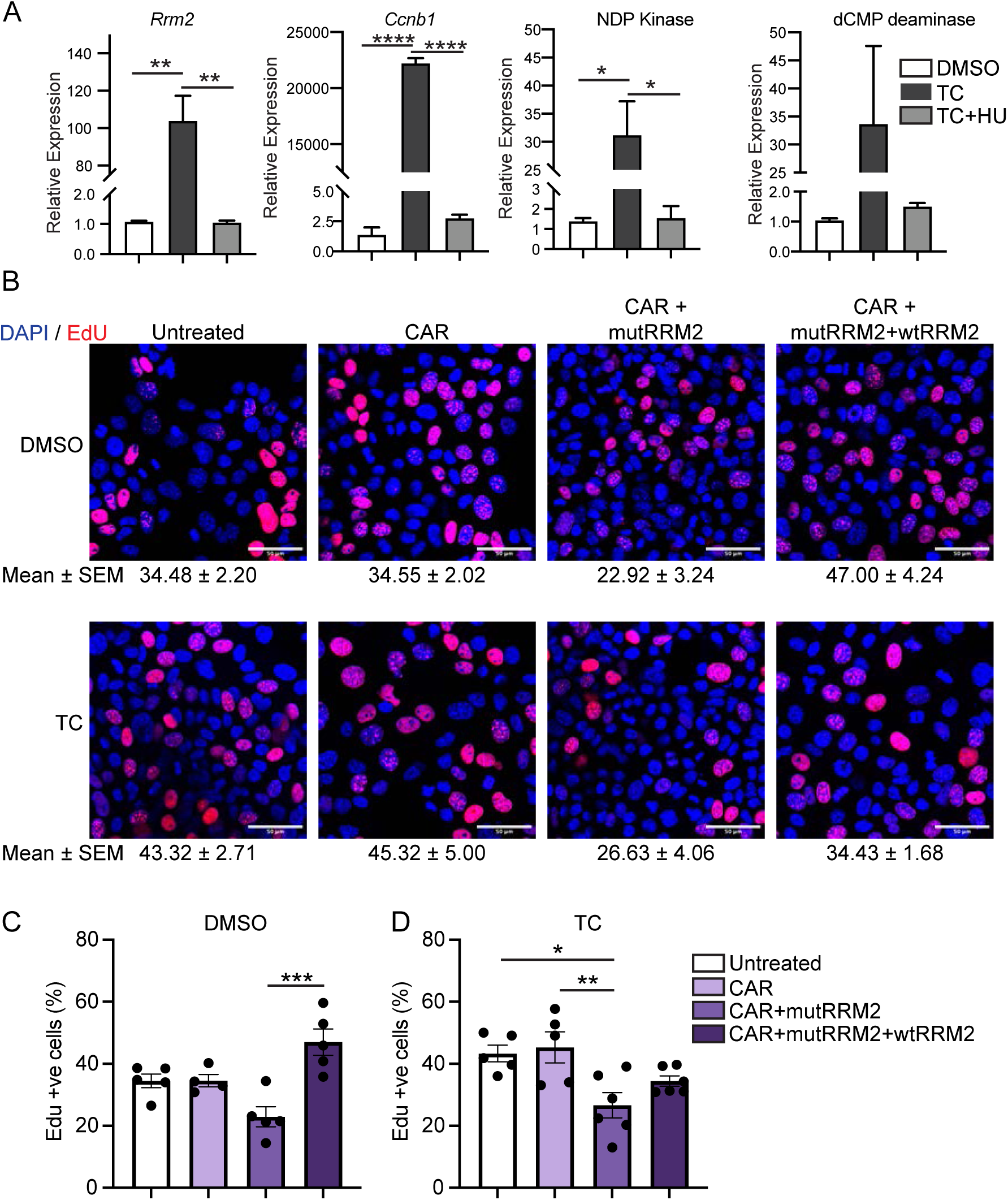
C**A**R**-mediated DNA synthesis is dependent on catalytically active RRM2.** (A) Hepatocytes isolated from WT mice were treated with HU or TC, and the transcript levels of *Rrm2*, *Ccnb1*, NDP Kinase, and dCMP deaminase were measured by qPCR, with β-actin as the internal control (n=3 mice/group). One-way ANOVA was performed, and the graphs show mean ± SEM. (B) Representative images of Edu incorporated (Edu in red and DAPI in blue) into the AML12 cells transfected with the CAR, WT RRM2 (wtRRM2), and Y176F mutant RRM2 (mutRRM2) constructs. The top panel corresponds to the DMSO, while the bottom panel corresponds to the TC treatments. (C-D) Quantification of Edu positive cells to the total number of cells under DMSO (C) and TC (D) treatment. One-way ANOVA was performed, and graphs are presented as mean±SEM.

Although wtRRM2 supplementation to mutRRM2+CAR revealed higher EdU incorporation, unlike the basal conditions, it did not fully compensate for it (Fig 5 B-D, Supplementary Fig 9 A-C). Taken together, these data provide compelling evidence that catalytically inactive RRM2 can dampen CAR-mediated DNA synthesis.

## Discussion

In rodents, transient CAR activation results in hepatomegaly, increased Ki-67 staining, and is linked to higher polyploidy, while prolonged activation results in liver cancer(14, 44). Although *FoxM1, Mdm2,* and *cMyc* signaling are implicated in facilitating CAR-driven liver proliferation, how CAR orchestrates DNA synthesis and ploidy remains unknown. Here, we identified a new CAR target, *Rrm2*, which encodes the catalytic subunit of the ribonucleotide reductase complex that is essential for *de novo* dNTP synthesis. Additionally, ligand activation of CAR was sufficient to induce several genes involved in dNTP biosynthesis. All these effects were absent in CARKO mice. We have not investigated if all these dNTP biosynthesis genes are direct or indirect targets of CAR. But one of the key rate-limiting steps is catalyzed by ribonucleotide reductase, and we demonstrate that CAR directly regulates the RRM2 subunit.

With respect to ploidy, CAR has been thought to increase ploidy by facilitating the endoreduplication of DNA(14). Here, we show that CAR can regulate various aspects of hepatic polyploidization. We found that CAR activation induces the *E2f* family of transcription factors, for instance, *E2f1* promotes polyploidization while *E2f*8 inhibits it. Moreover, we find that cytokinesis mediators- *Aurkb* and *Plk1* transcripts are also induced in a CAR-dependent manner. Thus, CAR may transcriptionally control several aspects of liver polyploidization and may contribute to maintaining overall ploidy distribution.

We validated that dNTP levels in livers were higher post-CAR activation, which is consistent with the *Rrm2* gene and protein induction. We find a significant increase in dATP and dTTP levels. dATP, in turn, can act as an allosteric inhibitor of RRM2 activity(36). Thus, RRM2 expression and activity are tightly controlled. One possibility is that the RRM2 increase seen in TC treatment is secondary to cell cycle activation to promote DNA synthesis. So, we tested the consequence of OE of RRM2 in quiescent CARKO livers from adult mice. We note that control viral treatments, as expected, result in a mild pro-proliferative response in both genotypes. While RRM2 OE in WT mice led to Ki-67 staining comparable to that of GFP virus-treated mice, in CARKO, it robustly amplified Ki-67 staining, indicating increased DNA synthesis in CARKO. PCNA also showed a similar pattern to Ki-67, except for higher basal staining with GFP virus-in CARKO. This could be due to PCNA staining not being restricted to the S phase. Therefore, the increased PCNA and Ki-67 staining in CARKO suggests that RRM2OE is sufficient to increase DNA synthesis, change ploidy, and affect the overall liver status size. Since other RRM subunits, especially *Rrm2b*, exhibit compensatory effects, we measured *Rrm1* and *Rrm2b* and found their levels unchanged upon RRM2 OE (Fig 4B, Supplementary Fig 4E).

A limitation of this study is that we do not fully understand how RRM2 OE promotes this heightened expression of S-phase markers in CARKOs. First, we examined the transcripts involved in the dNTP synthesis pathway. Post RRM2 OE, there was no change except for a reduction in UMP synthetase and dTMP kinase transcripts, indicative of a potential compensatory effect (Supplementary Fig 8 A-F). Similarly, cell cycle pathway genes were examined, and we found that S-phase cyclins (*Ccna1* and *Ccne1*) were increased in CARKO (Supplementary Fig. 8 G-H), which is consistent with the higher Ki-67 staining. However, the *E2f1* and *E2f8* transcripts did not change post-RRM2 OE in WT and CARKO livers (Supplementary Fig 8 I-J). These findings indicate that the ectopic introduction of RRM2 is sufficient to alter genes involved in the *de novo* dNTP biosynthesis pathway and S-phase cyclins, mainly in CARKO.

In terms of ploidy, RRM2 OE in the absence of CAR increased diploid nuclear content that coincided with the decrease in tetraploid nuclear content (Supplementary Fig 7 B-C). The cellular ploidy of CARKO RRM2 OE mimicked that of WT RRM2 OE, suggesting the CAR-mediated RRM2 expression is essential in maintaining the liver ploidy status (Fig 4 K-N). Polyploid hepatocytes have been shown to have a distinct metabolic capacity(7, 8, 45) and are in line with the xenosensing function of CAR.

We demonstrate that CAR promotes DNA synthesis directly, as measured by EdU incorporation, and that this increase is contingent on the presence of a functional RRM2, as transfection of a catalytically inactive RRM2 quenched EdU labeling observed upon co-transfection of CAR and wtRRM2. Overall, these data clearly show that CAR-RRM2 signaling may coordinate liver proliferation and polyploidy status and that CAR may play a basal role in maintaining liver ploidy status during development.

## Data Availability

All data produced in this study are included in the article and in its online supplementary material.

## Acknowledgements

We would like to thank Dr Chang Cui, Department of Biochemistry, UIUC, for helping in critically interpreting RRM2 overexpression data and for sharing RRM2 constructs and suggesting appropriate catalytically inactive RRM2 mutants. We also thank Dr. Auinash Kalsotra, Department of Biochemistry, UIUC, for valuable advice, resources and inputs throughout this project. Core Facilities at the Carl R. Woese Institute for Genomic Biology, at the UIUC and at The University of Pittsburgh Unified Flow Core (RRID:SCR_025102) supported this project.

## Authors Contribution

S.A conceived, conceptualized and obtained funding for this study. A.A., B.M., and S.A. planned and designed the experiments for this study. D.D.M was involved in the initial study design. S.A performed the initial protein, RNA, ChIP and luciferase analysis. B.M. and A.C. performed the *de novo* dNTP pathway analysis. L.W. and C.K.M. performed the dNTP analysis. F.A. and A.W.D. performed all the FACS sorting and ploidy data analysis. B.M., A.A. and B.C. isolated hepatocytes and nuclei for FACS analysis. A.A. and S.A. designed and carried out the experiments for overexpression studies. B.M, and A.C performed RRM2 inhibitor studies. S.C. did all the transfections for EdU staining and S.C and A.A imaged and analyzed the data. A.A., A.W.D and S.A. wrote the manuscript with suggested edits by all the authors.

