## Supplementary Figures for "Constitutive Androstane Receptor induces Ribonucleotide Reductase-M2 expression and maintains hepatocyte ploidy in mice"

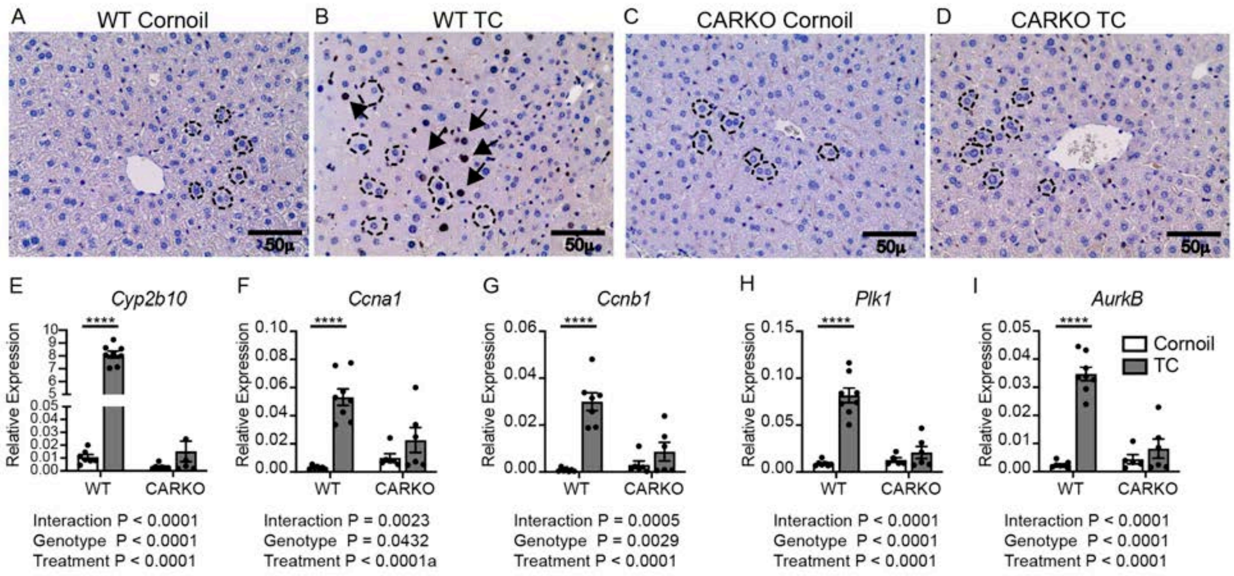


**Supplementary Fig 1 – CAR activation leads to an increase in Cyclins and mitotic entry genes.** (A-D) Representative images of DNA synthesis marker, Ki-67 staining for WT (A and B) and CARKO (C and D) with and without TC treatment. The black arrow points to the Ki-67 staining and dotted line markers of the hepatocyte cell boundary. (E) CAR activation on TC treatment was determined by examining *Cyp2b10* transcript levels. (F-I) The Transcript levels of cyclins (*Ccna1*) and *Ccnb1*) and mitotic kinases (*Plk1* and *AurkB*) genes were analyzed using qPCR (n = 6-8 mice/group, adult male mice, 3-month-old, 36B4 was used as the internal control).

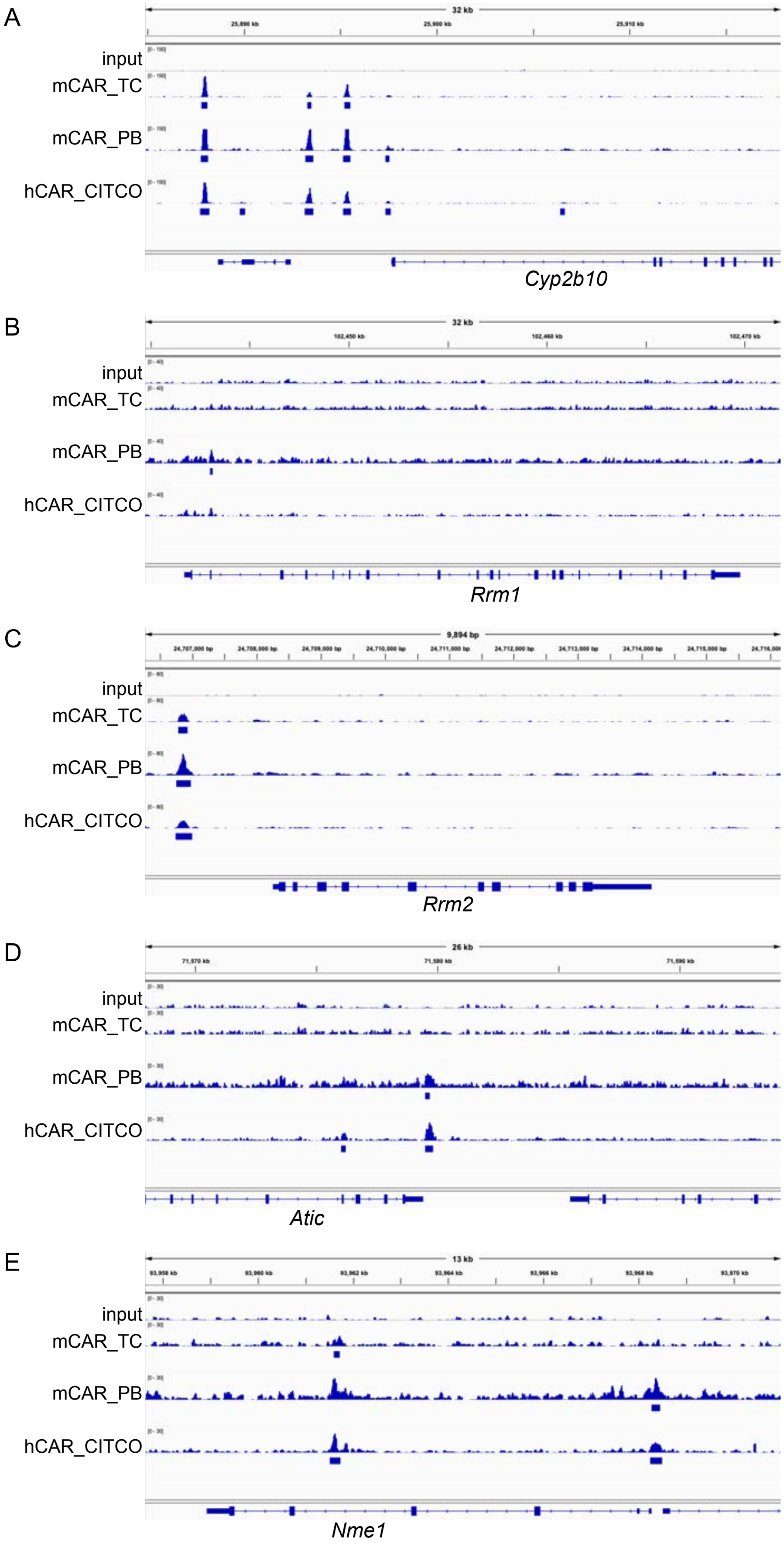
A
32 kb
input
mCAR_TC
mCAR_PB
hCAR_CITCO
Cyp2b10
B
32 kb
input
mCAR_TC
mCAR_PB
hCAR_CITCO
Rrm1
C
9,894 bp
input
mCAR_TC
mCAR_PB
hCAR_CITCO
Rrm2
D
26 kb
input
mCAR_TC
mCAR_PB
hCAR_CITCO
Atic
E
13 kb
input
mCAR_TC
mCAR_PB
hCAR_CITCO
Nme1

**Supplementary Fig 2 – ChIP sequence analysis of CARKO mice expressing YFP tagged mCAR or hCAR and treated with PB, TC or CITCO (known CAR agonist)**. Previously published CAR-ChIP seq data was mined and we found enrichment of CAR binding to the its well-known target gene (A) *Cyp2b10*, (B) *Rrm1*, (C) *RRM2,* (D) *Atic*, and (E) *Nme1* (encoding for NDP kinase). Thus, several involved in *de novo* dNTP synthesis pathway exhibit potential CAR-mediated regulation.

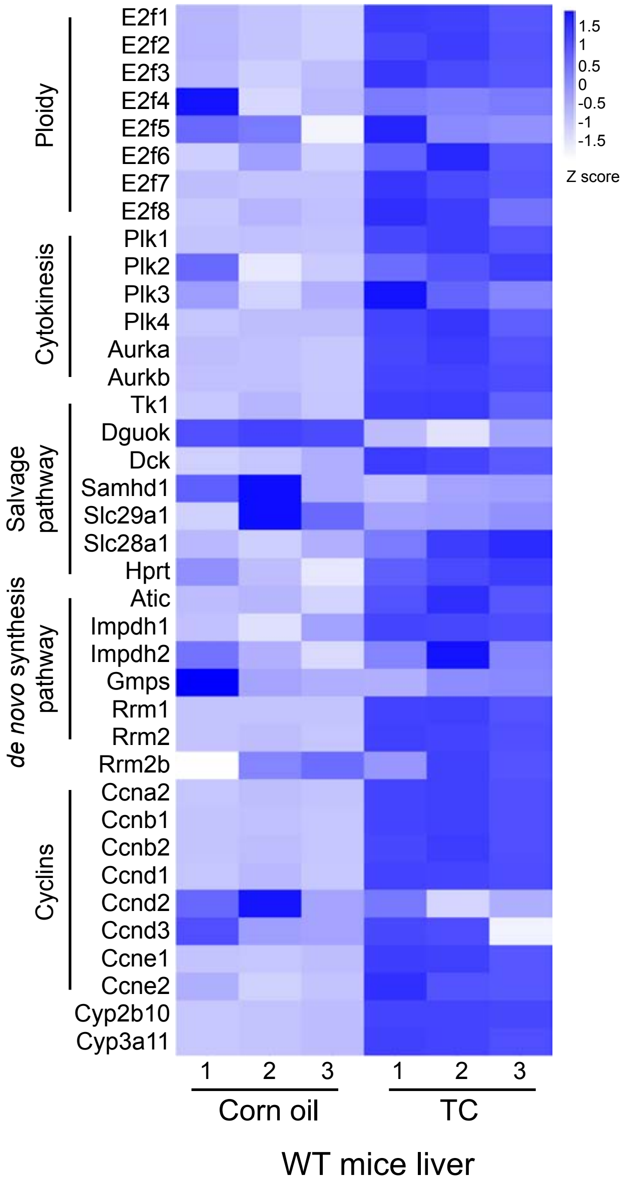


**Supplementary Fig 3 – Pro-proliferative gene expression on TC treatment**. RNA seq analysis of WT mice with Cornoil or TC treatment[24] shows that, besides the anticipated increase in Cyp genes and cyclins, we also found induction in the dNTP salvage pathway, *de novo dNTP* synthesis pathway, cytokinesis, and ploidy-regulating genes upon TC treatment.

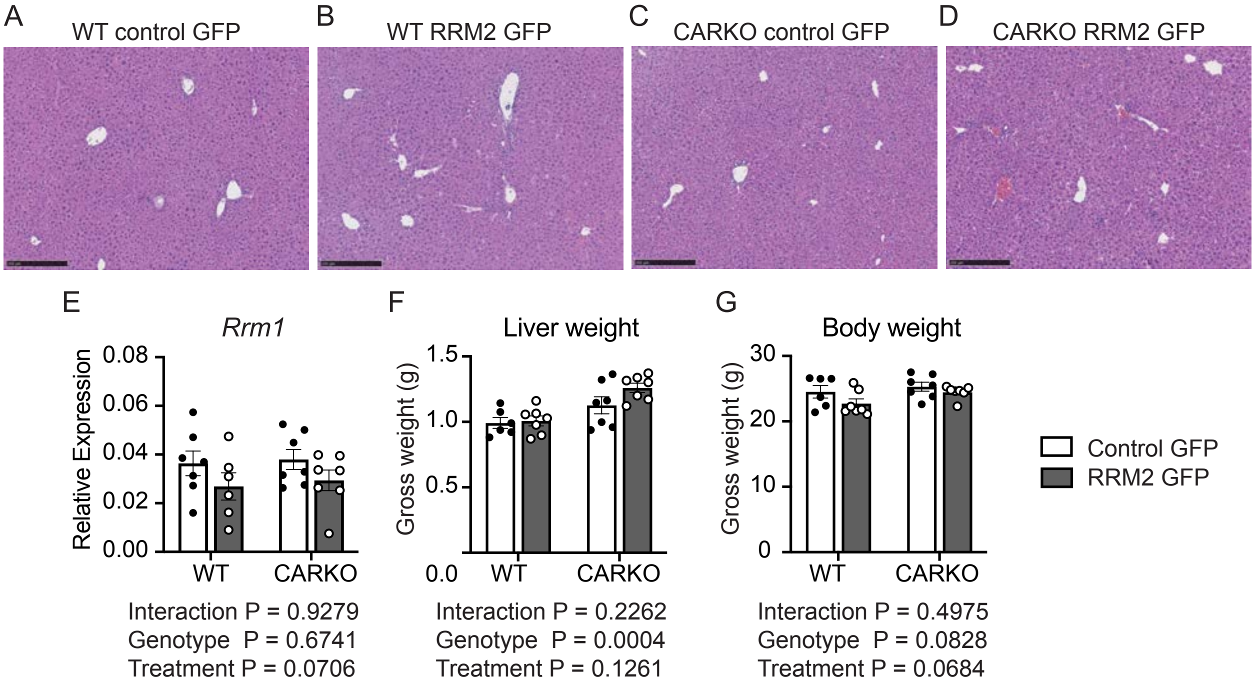


**Supplementary Fig 4 – AAV-based RRM2 overexpression did not induce inflammation or expression of other subunits of ribonucleotide reductase**. (A-D) Representative H&E staining of WT and CARKO mice liver 2 weeks post-treatment with control GFP or RRM2-GFP. (E) *Rrm1* transcript levels were analyzed from WT and CARKO mice livers under overexpression of RRM2 using qPCR. (n = 6-8 mice/group, 4-month-old female mice, 36b4 was used as the internal control for qPCR analysis). (F) Gross liver and (G) body weight for the viral-treated mice.

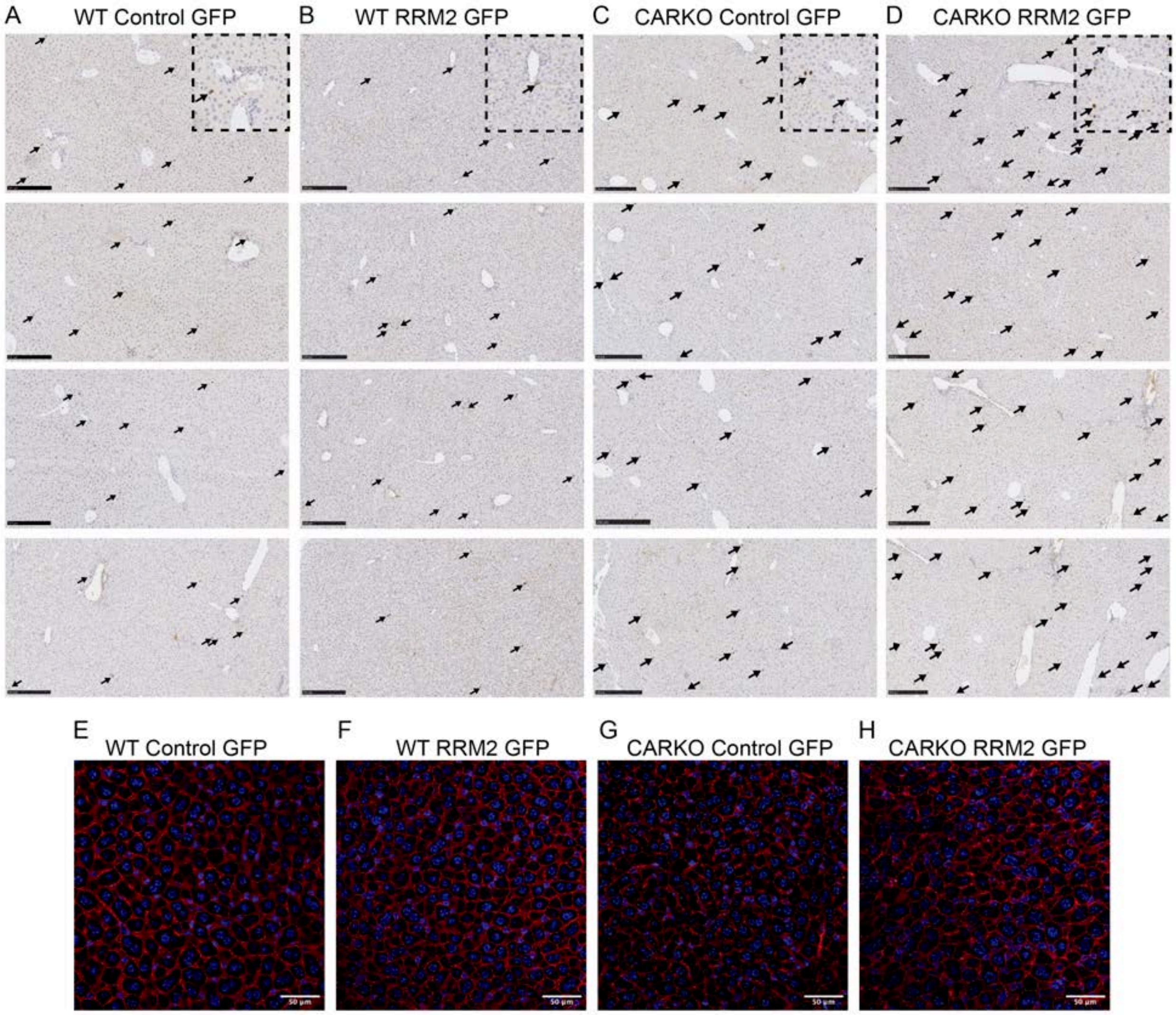


**Supplementary Fig 5 – Ki-67 and β-catenin staining for WT and CARKO liver with RRM2 overexpression**. (A-D) Representative Ki-67 images of WT (A-B) AND CARKO (C-D) mice treated with Control-GFP (A and C) or RRM2-GFP (B and D) virus. Black arrows point to the Ki-67 stained using Immunohistochemistry. Enlarged images of Ki-67 staining are presented as insets in the upper panel of the figures in the right corner. (E-H) Representative images showing DAPI and β-catenin staining in control-GFP and RRM2-GFP for WT and CARKO mice livers. The hepatocyte area was measured using ImageJ software. (n = 6-10 mice/group)

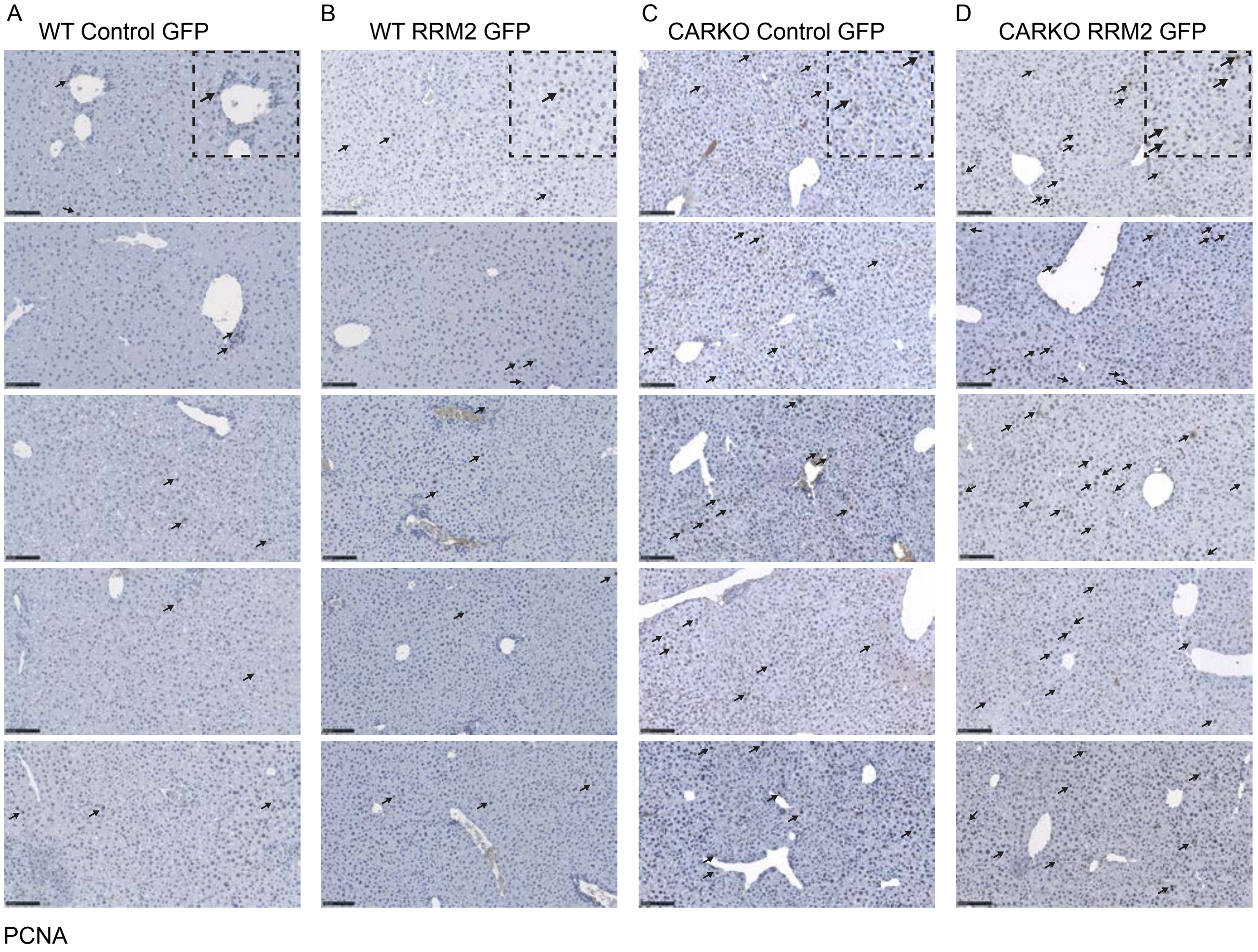


**Supplementary Fig 6 – PCNA staining for WT and CARKO liver with RRM2 overexpression**. (A-D) Representative PCNA images of WT (A-B) AND CARKO (C-D) mice treated with Control-GFP (A and C) or RRM2-GFP (B and D) virus. Black arrows point to the PCNA-stained hepatocytes using Immunohistochemistry. Enlarged images of PCNA staining are presented as insets in the upper panel of the figures on the right corner. (n = 3-4 mice/group)

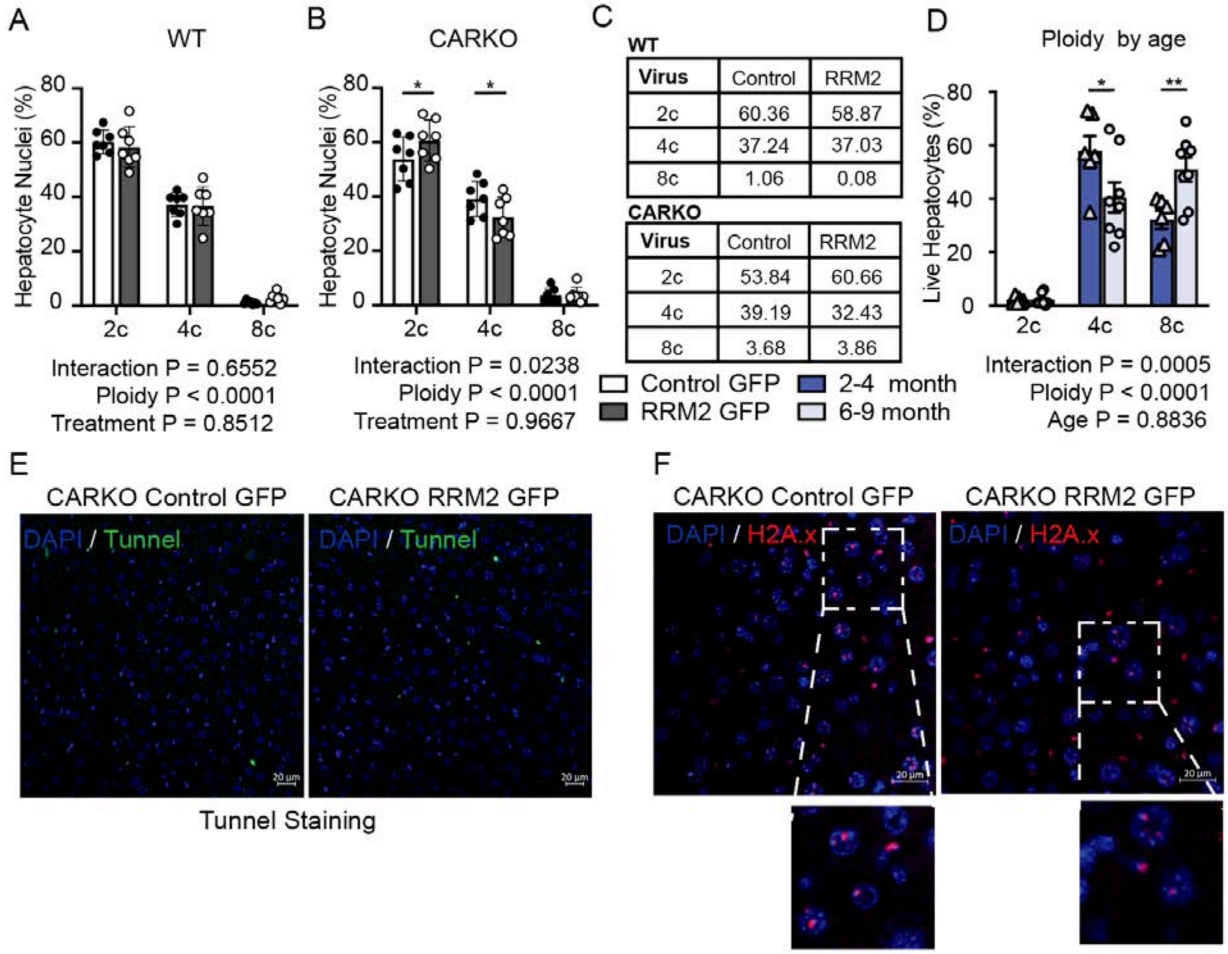


**Supplementary Fig 7– Overexpression of RRM2 rebuilds the lost ploidy in CARKO hepatocytes to the WT condition**. (A-B) Quantification for the liver nuclear ploidy analysis of WT and CARKO mice under RRM2 overexpression, respectively (same sample set used for the gene expression analysis to prevent batch error between the treatment groups, n = 7 mice/treatment group). (C) Average nuclear ploidy analysis across 5-6 sample sets. (D) Age-dependent cellular ploidy analysis from WT male mice. (n=6-8 mice/age). (E) Representative images of tunnel staining in the CARKO control and RRM2 OE. (F) Representative images of H2A.x staining between the control and RRM2 OE virus in CARKO mice.

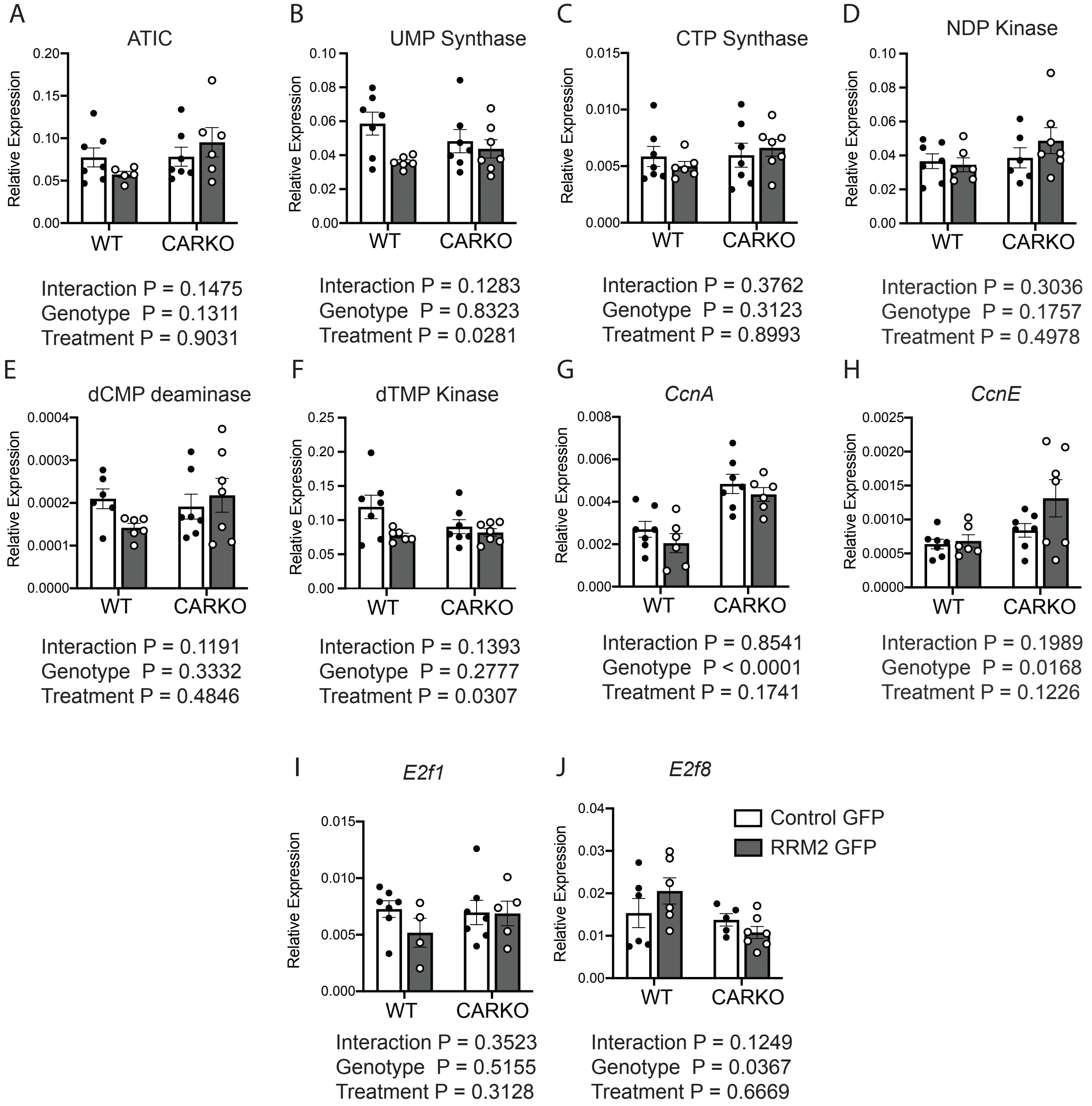


**Supplementary Fig 8 - No change in the enzyme and S and G2 cyclin transcript levels OR E2Fs after 2 weeks of RRM2 OE**. qPCR analysis of A) ATIC, B) UMP synthase, C) CTP synthase, D) NDP kinase, E) dCMP deaminase, F) dTMP Kinase, G) Cyclin A (*Ccna*), H) Cyclin E (*Ccne1*), (I) *E2f1,* and (J) *E2f8,* normalized to the internal control 36B4. There was no significant difference between the Control and RRM2 GFP treatment in WT or CARKO mice models.

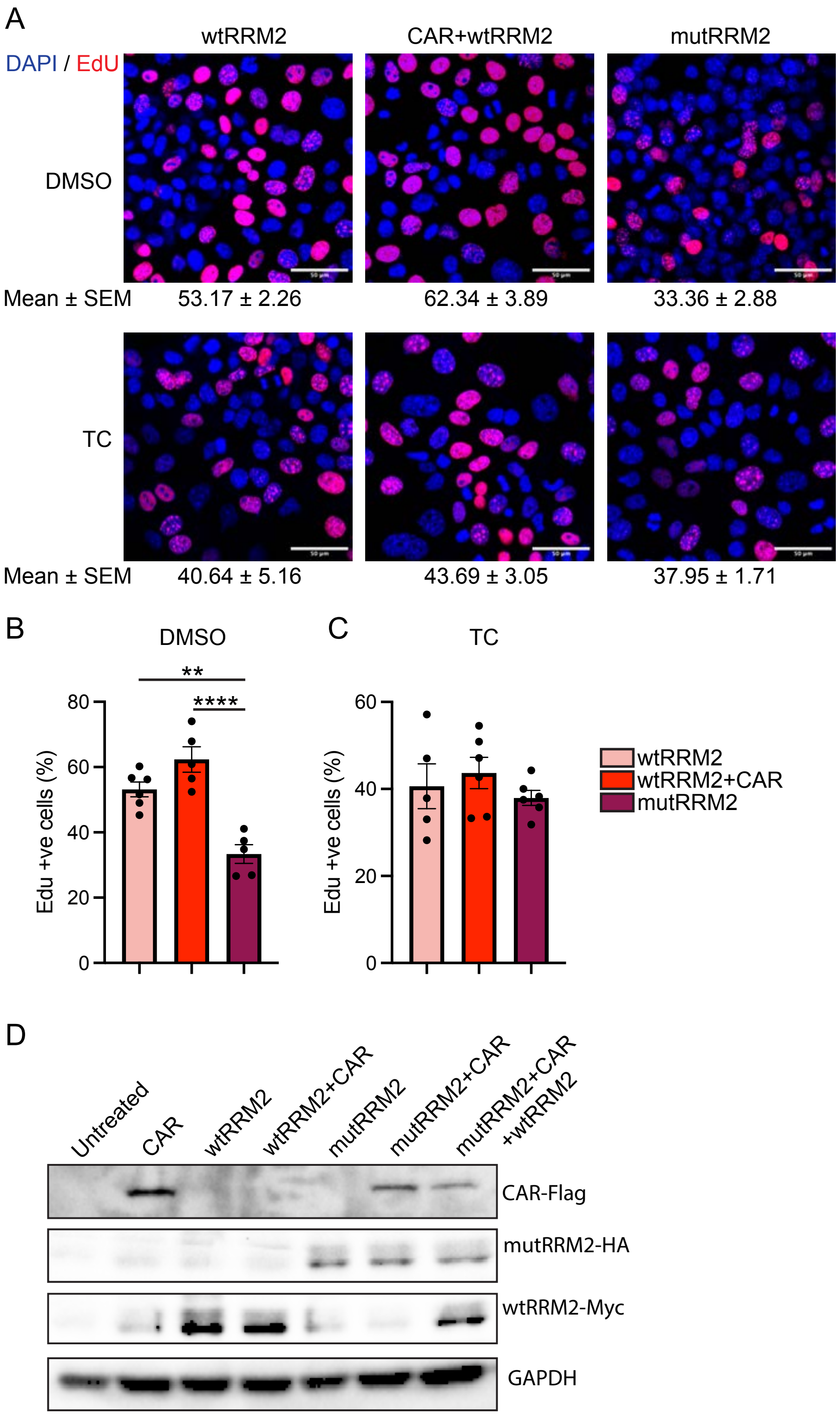


**Supplementary Fig 9 - WT RRM2 along with CAR co-transfection promotes proliferation.** (A) Representative images of Edu incorporated (Edu in red and DAPI in blue) into the AML12 cells transfected with wtRRM2, CAR, and mutRRM2. The top panel corresponds to the DMSO, while the bottom panel corresponds to the TC treatments. (B-C) Quantification of Edu positive cells to the total number of cells under DMSO (B) and TC (C) treatment. One-way ANOVA was performed, and graphs are represented as mean$\pm$SEM. (D) Immunoblot from AML12 cell lysate prior to TC or DMSO treatment indicating similar level of individual construct transfection between different group.

**Supplementary Table 1 – List of primers used in this study**

| Gene | Forward Primer (5’-3') | Reverse Primer (5’-3’) |
|---|---|---|
| *36B4* | AGATGCAGCAGATCCGCAT | GTTCTTGCCCATCAGCACC |
| β-*actin* | TCTCCAGGGAGGAAGAGGAT | GCTACAGCTTCACCACCACA |
| *Rrm1* | ATGTGATCAAGCGAGATGGC | TTCATGGTGATCTGAGCAGG |
| *Rrm2* | GTTGTCTTTCCCATCGAGTACC | GAGCTTCCCAGTGCTGAATATC |
| *Rrm2b* | GAGCCACTCCTAAGAAAGAGTTC | GAGGGAGGTCCTTTGACAAGT |
| *Cyp2b10* | TGCTGTCGTTGAGCCAACC | CCACTAAACATTGGGCTTCCT |
| *Ccna1* | AGAAGACAAGCCAGTGAACG | GTCTGGTTGCCTCTTCATGT |
| *Ccnb1* | GCCAAGAGCCATGTGACTATC | CAGAGCTGGTACTTTGGTGTTC |
| *Ccne1* | GATCGTTACATGGCATCACA | AAACTGGTGCAACTTTGGAG |
| *E2f1* | ATCACCTCCCTCCACATCC | TGACAGTTGGTCCTCTTCCA |
| *E2f8* | GCAGTTGGAAGAGCAGTCAA | GCTGGATAAGGGACGGTGTA |
| *Plk1* | TTTGAGGTGGATGTGTGGTC | ACTGGGTTGATGTGCTTGG |
| *Aurkb* | GGATGTTGGGATGTTTCAGG | GGAGAAGAAGAGCCGTTTCA |
| ATIC | TGTGGTCTGTAACCTGTACCC | CACGCCACCAATATCAATTTGTT |
| UMP | CAGTCAGGTCGCAGACATTTT | CCAGTGGTAACGCTGTATAAGGA |
| CTP | CTTTCTCCAGACTTAGTGGTGTG | ACCCGGTAGATGGATGAAACA |
| NDP | GAGGACTGCCGGAGGTTTTAC | GCAAGGATATAGGCTCGGATTG |
| dCMP | TCGAAGGAGCGGACGTGA | CTGACAAGAAGGCCACTGC |
| dTMP | TGCGTTTCCCCGAAAGATCAA | CCCAGCGGTTTGCAGAGAA |
